# Prevalence of errors in lab-made plasmids across the globe

**DOI:** 10.1101/2024.06.17.596931

**Authors:** Xingjian Bai, Jack F. Hong, Shan Yu, David Y. Hu, Amy Y. Chen, Constance A. Rich, Silk J. Shi, Sandy Y. Xu, Daniel M. Croucher, Kristofer J. Müssar, Daniel W. Meng, Jane L. Chen, Bruce T. Lahn

## Abstract

Plasmids are indispensable in life sciences research and therapeutics development. Currently, most labs custom-build their plasmids. As yet, no systematic data on the quality of lab-made plasmids exist. Here, we report a broad survey of plasmids from hundreds of academic and industrial labs worldwide. We show that nearly half of them contained design and/or sequence errors. For transfer plasmids used in making AAV vectors, which are widely used in gene therapy, about 40% carried mutations in the inverted terminal repeat (ITR) regions due to their inherent instability, which is influenced by flanking GC content. We also list genes difficult to clone into plasmid or package into virus due to their toxicity. Our finding raises serious concerns over the trustworthiness of lab-made plasmids, which parallels the underappreciated mycoplasma contamination and misidentified mammalian cell lines reported previously, and highlights the need for community-wide standards to uphold the quality of this ubiquitous reagent in research and medicine. Accordingly, we propose the concept of good vector practice (GVP) that covers the proper design, construction, in-process QC, final QC, banking and management of plasmids in research and medicine to uphold their quality.

## Introduction

Plasmids are extrachromosomal DNA capable of independent replication in cells. They are most commonly found in bacteria in circular double-stranded form. Plasmids were first identified in bacterial antibiotic resistance studies in the 1950s (1). In the 1970s, recombinant DNA technology enabled the engineering of artificial plasmids carrying foreign DNA of interest (2), which in the ensuing years propelled plasmids to become a central and ubiquitous reagent in the life sciences. Nowadays, plasmids are used mostly as gene delivery vectors in vitro and in vivo, either directly or as starting materials for generating viral and mRNA vectors. In addition to their ubiquity in research, plasmids have also become a foundational source material in manufacturing many therapeutic products such as recombinant protein drugs including antibodies, gene therapy vectors, and the recent COVID-19 mRNA vaccines.

Plasmids are a highly customized reagent because different experimental applications generally require different plasmids. For many decades, researchers have typically constructed their own plasmids in the lab or shared them from other researchers. As yet, there is no systematic quality assessment of lab-made plasmids on a global scale despite their importance in research and medicine, likely because such an endeavor would be impractical for any single lab.

As a cloning service provider, we received a large number of lab-made plasmids along with their theoretical sequences from both academia and industry across the world, which accorded us an opportunity to systematically assess their quality. We observed a wide variety of design errors ranging from obvious ones that most trained molecular biologists can identify, to subtle mistakes that even very seasoned experts may not spot. Sequence errors are even more prevalent. In combination, design and sequence errors affect nearly half of the lab-made plasmids we received.

We paid special attention to AAV transfer plasmids used to package recombinant AAV virus because they were often used to develop gene therapy drugs. We found that their ITRs were highly mutable, with about 40% of the plasmids we received bearing mutations relative to wildtype ITR sequence. We further showed that ITR instability is associated with high GC content of the immediate flanking sequence.

Researchers typically send plasmids to us for further sequence modification, recombinant viral vector production, and/or in vitro and in vivo experiments. Given that the senders are devoting significant financial resources and time to contract us to perform these downstream projects, they have a vested interest in ensuring the correctness of their plasmids. Considering this, it is possible that the quality issues we uncovered might underestimate the true scale of the problems in labs. Similar to the reports of mycoplasma contamination and misidentified mammalian cell lines (3-5), our comprehensive survey shines a spotlight on significant quality issues with lab-made and shared plasmids in academia and industry around the world.

## Materials and methods

### Sample collection

Our global clients submitted their lab-made plasmids to be modified, used as cloning materials, packaged into recombinant viruses, employed as templates for making RNA by in vitro transcription (IVT), or used in other molecular biology services. These starting materials were required to be submitted as DNA solution (>1 μg dissolved in 0.1-1 X TE) or bacterial stab culture, along with their theoretical vector maps and sequences. For AAV transfer plasmids used for the packaging of recombinant AAV virus, the two ITRs flanking the payload region are in theory identical in sequence and are therefore indistinguishable based on sequence alone. By convention, we refer to the ITR closer to the replication origin on the plasmid backbone as the 5’ ITR (aka left ITR or upstream ITR), and the other ITR as the 3’ ITR (aka right ITR or downstream ITR). Some of our clients labeled their ITRs in the opposite way, which we changed to the above standard convention for consistency.

### Sample analyses

Before project initiation, the submitted plasmids were subjected to our standard quality control (QC) protocols. The designs of the plasmids were manually evaluated by our scientists to ensure their correctness for the intended applications. The structures and sequences of the plasmids were validated by restriction digestion and/or Sanger sequencing. Comparison between Sanger sequencing and other commonly used sequencing methods for validating plasmids such as Nanopore sequencing showed that in all cases where discrepant results were produced, Sanger always gave the correct sequences. Additionally, the error rate of Sanger sequencing is the lowest among all sequencing methods based on our experience and the literature. We therefore relied exclusively on Sanger. Indeed, we recommend that Sanger be considered by the community as the gold standard for sequence validation of plasmids, especially in cases where sequence accuracy of plasmids is of the utmost importance such as in clinical applications. Two well-tested restriction enzymes with theoretical cut sites on the plasmid map that should generate bands with distinguishable sizes were applied together or individually on the plasmid DNA and analyzed with gel electrophoresis. Bands produced were compared to theoretical bands from the submitted plasmid maps and sequences. Regions of the vector critical to biological functions or downstream applications, such as cloning, virus packaging, and IVT RNA production (e.g. cloning sites, coding sequences, promoters, UTRs, PolyA signal sequences, lentivirus LTRs, AAV ITR, etc.), were validated by Sanger sequencing. Due to confidentiality obligations to our clients, we are unable to provide detailed maps and sequences of their original plasmids here. However, to better showcase design errors without breaching confidentiality, we added sample plasmids to the design errors listed in Table 1. The maps and sequences of these sample plasmids are viewable online through the URLs given in Table 1, but any confidential information related to our clients’ original plasmids are desensitized.

**Table 1.**
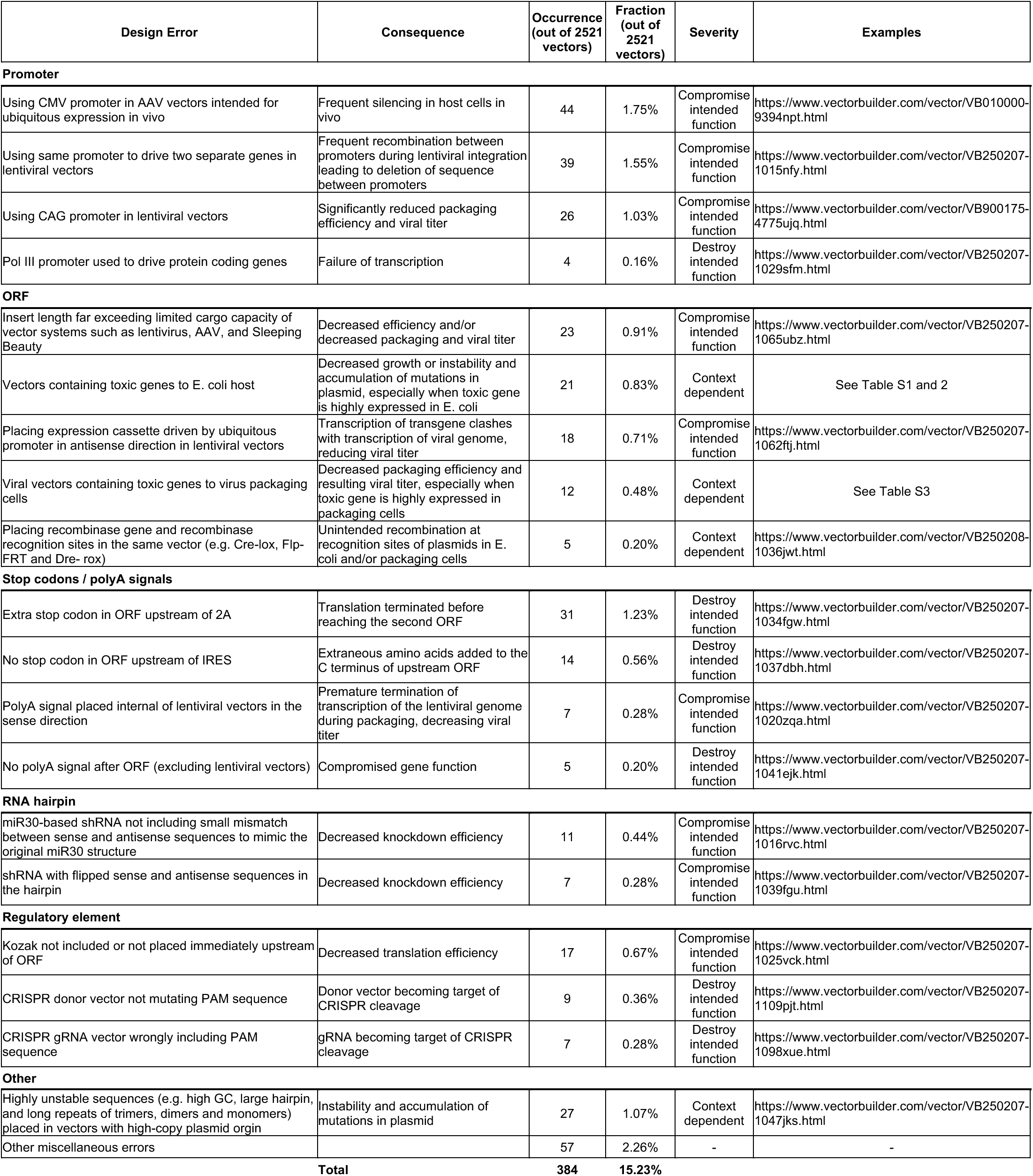
Design errors in lab-made plasmids.

#### Prevalence of design errors in lab-made plasmids

When receiving plasmids from researchers to perform various projects, such as virus packaging and IVT RNA production, or to clone another vector they designed, we would first evaluate the reference or the final maps provided by the senders to ensure that the vectors were appropriately designed. Strikingly, of 2,521 plasmids that we received in a certain period from academia and industry around the world, we found that about 15% (384) contained significant design errors that could impact function. Design errors were found in most types of components, with the most prevalent being the incorrect choice or placements of promoters, followed by problems in the choice or design of open reading frames (ORFs). While certain design errors might serve specific purposes intended by the researchers, we note that nearly all researchers accepted our recommended corrections of the errors. Details of common design errors and their corresponding consequences and frequencies are listed in Table 1. We also rated these errors by levels of significance that reflect their impact on function, including “Destroy intended function” that destroys the intended function of the vectors, “Compromise intended function” that reduces the intended function to some degree, and “Context dependent” that may, under some circumstances, lead to destroyed or compromised function (Table 1). For each design error, we also provided a sample plasmid viewable online through the URL given to illustrate the nature of the error.

Many errors appear to be due to limited understanding of the nuances in designing appropriate gene delivery systems, such as limits on cargo capacity, promoter silencing, and considerations for different types of linkers (Table 1). Other problems are related to the biological properties of specific sequences in individual vectors. Some sequences are unstable in *E. coli*, such as long inverted repeats that are prone to form large hairpins, extremely high-GC sequences, and short tandem repeats (e.g. long strings of mono-, di-, or tri-nucleotides, including A tracts in template plasmids for making IVT RNA). When these sequences are cloned into high-copy plasmids, they can quickly accumulate mutations, including large deletions and rearrangements (Table 1). For greater stability, it is necessary to clone these sequences into low-copy plasmids along with tailored *E. coli* hosts, and utilize special culture conditions such as low temperature, low salt, and adjusted antibiotic concentration. The sequences themselves can often be modified to increase stability while maintaining intended biological functions, e.g. placing a short intervening sequence in the long A tract in in vitro transcription plasmids.

Additionally, a major issue that sometimes plagues lab-made plasmids is toxicity of the gene of interest (GOI) that they carry. If the GOI is toxic to *E. coli*, then cloning it into a plasmid can be very difficult and sometimes impossible. In cases where cloning is successful, the GOI or its surrounding sequences tend to be highly unstable and can quickly accumulate mutations that compromise GOI function (unpublished data), presumably due to strong selective pressure against the intact toxic form of the gene. By design, there are typically no prokaryotic promoters driving expression of the toxic GOIs, so the fact that they can still exert their detrimental effect on the host indicates the presence of cryptic promoters driving their expression in *E. coli*.

Similarly, genes contained in viral transfer plasmids can be toxic to packaging cells or interfere with virus packaging pathways, leading to dramatically reduced packaging efficiency and viral titer. Unfortunately, toxicity of genes is often hard to predict even for labs working with them.

Table S1 and S2 list genes showing toxicity to *E. coli* host that we have encountered. Table S1 contains 46 genes that are moderately toxic, and their cloning in intact forms can often be accomplished by employing various workarounds such as using low-copy plasmids (such as pColE1 and pSC101), switching to different *E. coli* host strains, and altering culture conditions. Table S2 lists 25 genes that are severely toxic, and their cloning in intact forms was unsuccessful in our hands by the above workarounds alone, though we managed to clone most of them in various mutated forms such as introducing point mutations or truncations, and inserting synthetic introns. These toxic genes are enriched for membrane channels and transporters, and proteins involved in DNA dynamics such as DNA repair, topoisomerase activity, and chromosome segregation. Particularly striking is the enrichment for calcium and sodium channels, with each type accounting for about ten (14%) of the toxic genes listed. There are also two chloride channels on the list. This enrichment is presumably due to these channel genes causing ion imbalances in *E. coli* host. Indeed, the toxicity of ion channel genes in cloning may be a rule rather than an exception.

Table S3 lists 73 genes from specific species that we found to be toxic to virus packaging, resulting in very low viral titer in at least some cases. They are enriched for pro-apoptotic genes (e.g. BAX and N-GSDME), cell cycle regulators (e.g. BABAM2 and NEK1), proliferation modulators (e.g. F2RL1 and Foxn1), and antiviral genes (e.g. EIF2AK2 and APOBEC3A). Interestingly, a gene that severely inhibits packaging efficiency when placed in a particular viral transfer plasmid may not have the same detrimental effect when placed in another transfer plasmid, presumably because its toxic effect also depends on other factors such as the strength of the promoter driving the GOI in packaging cells, the type of virus being packaged (e.g. lentivirus vs. AAV), and the packaging cell lines used. One solution often effective in reducing GOI toxicity to virus packaging is to use weaker or inducible promoters. For example, a lentiviral vector containing a medium-strength promoter driving mouse Foxn1 produced tenfold higher titer as compared to the same vector using a strong promoter.

To promote continuous sharing of experience, knowhow and encountered pitfalls in the design, construction and utilization of gene delivery vectors, we have committed to hosting an online Gene Delivery Forum where we and other researchers can share information about this topic. We have recently posted an article about the problem of using two identical promoters in a lentiviral vector and presented supporting data: https://www.vectorbuilder.com/resources/vector-academy/lessons/design-pitfall.html.

#### Prevalence of sequence errors in lab-made plasmids

We subjected 1,132 plasmids provided by researchers to further QC validation. Of these, about 1.9% (21/1132) could not be recovered from the *E. coli* stocks we received or the incorrect samples were sent. We analyzed the overall structure of 852 plasmids by restriction enzyme (RE) digestion, selecting multiple RE sites from the sender-provided vector maps and sequences that were expected to yield distinct fragments upon digestion. The other plasmids were sequenced directly without RE digestion. Remarkably, RE digestion of 852 plasmids revealed inconsistent fragment patterns in about 15% (128/852), indicating significant rearrangements of these plasmids or point mutations at the RE sites (Figure 1A). Given that RE digestion only confirmed the general structure of the plasmids, we also performed sequencing-based validation on some plasmids, focusing on functional regions utilized in downstream cloning or crucial for intended biological applications. Here, ITR regions of AAV transfer plasmids were excluded from analysis because their sequence mutations were evaluated separately (see detailed description below). Sanger sequencing was chosen over other sequencing methods such as next-generation sequencing (NGS) and third-generation (single molecule) sequencing such as Nanopore due to its much higher fidelity and reliability in validating plasmid sequences (see Materials and methods). We Sanger sequenced 117 plasmids with correct RE digestion patterns and found that about 24% (28/117) exhibited inconsistent sequences compared to the senders’ reference, and two failed to have their functional regions fully sequenced, presumably due to the presence of difficult sequences (Figure 1A). To remove any bias, we directly sequenced 259 plasmids without performing initial RE digestion, focusing on functional regions (again, excluding AAV ITRs). Notably, about 35% of these plasmids (91/259) displayed sequence variations from the senders’ reference (Figure 1B). Among them, we identified 89 point mutations, 35 deletions, and 19 insertions, with some plasmids containing multiple types of errors.

**Figure 1.**
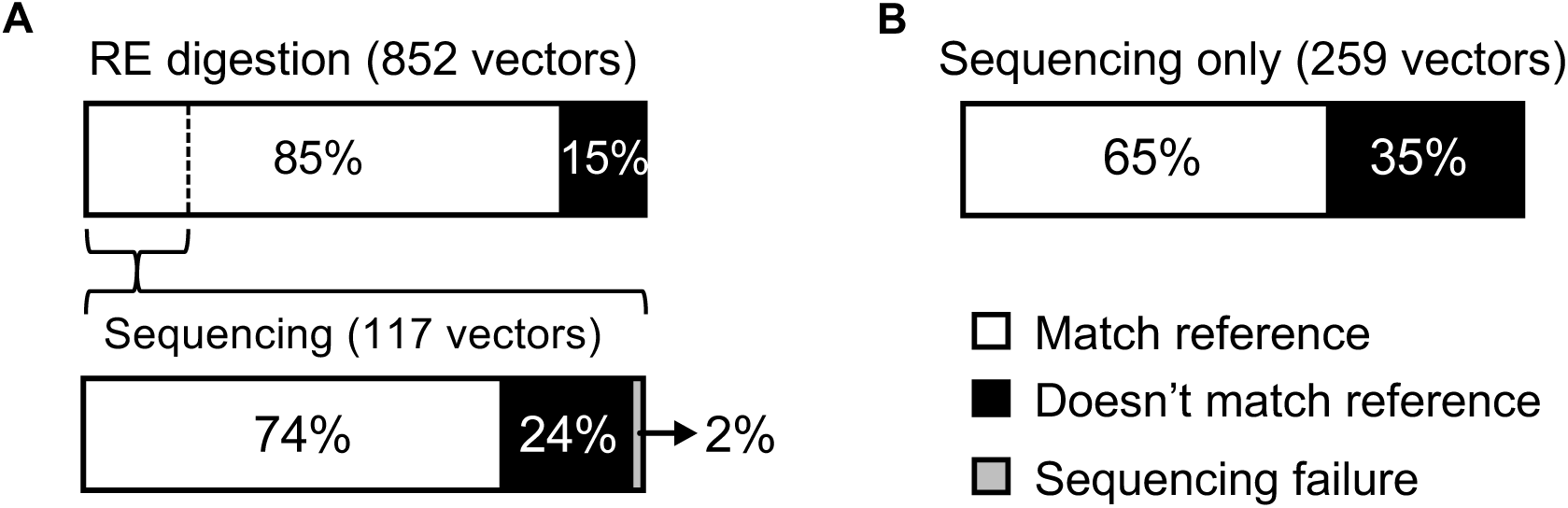
High rate of sequence errors in lab-made plasmids from global researchers. (**A**) Error rate of plasmids was assessed by restriction enzyme (RE) digestion, and a subset of RE-validated plasmids were further assessed by Sanger sequencing of their functional regions. (**B**) Error rate was directly assessed by Sanger sequencing of functional regions without RE digestion.

#### ITRs of AAV transfer plasmids are highly mutable

We paid special attention to AAV vectors given their therapeutic importance (6,7) A superior feature of AAV is that the only cis sequence elements required for packaging recombinant virus are the two ITRs flanking the payload sequence in the transfer plasmid (Figure 2A, 2B). By convention, ITRs of AAV serotype 2 (AAV2) are widely utilized in recombinant AAV vectors due to their compatibility with a wide range of capsid types (7-9). However, AAV2 ITRs contain over 70% GC and can form complex secondary structure. As a result, ITR sequences on AAV transfer plasmids can acquire mutations that impair packaging, leading to decreased full capsid ratio and increased encapsulation of cellular DNA, problems that can significantly compromise the use of recombinant AAV as a therapeutic agent (10,11).

**Figure 2.**
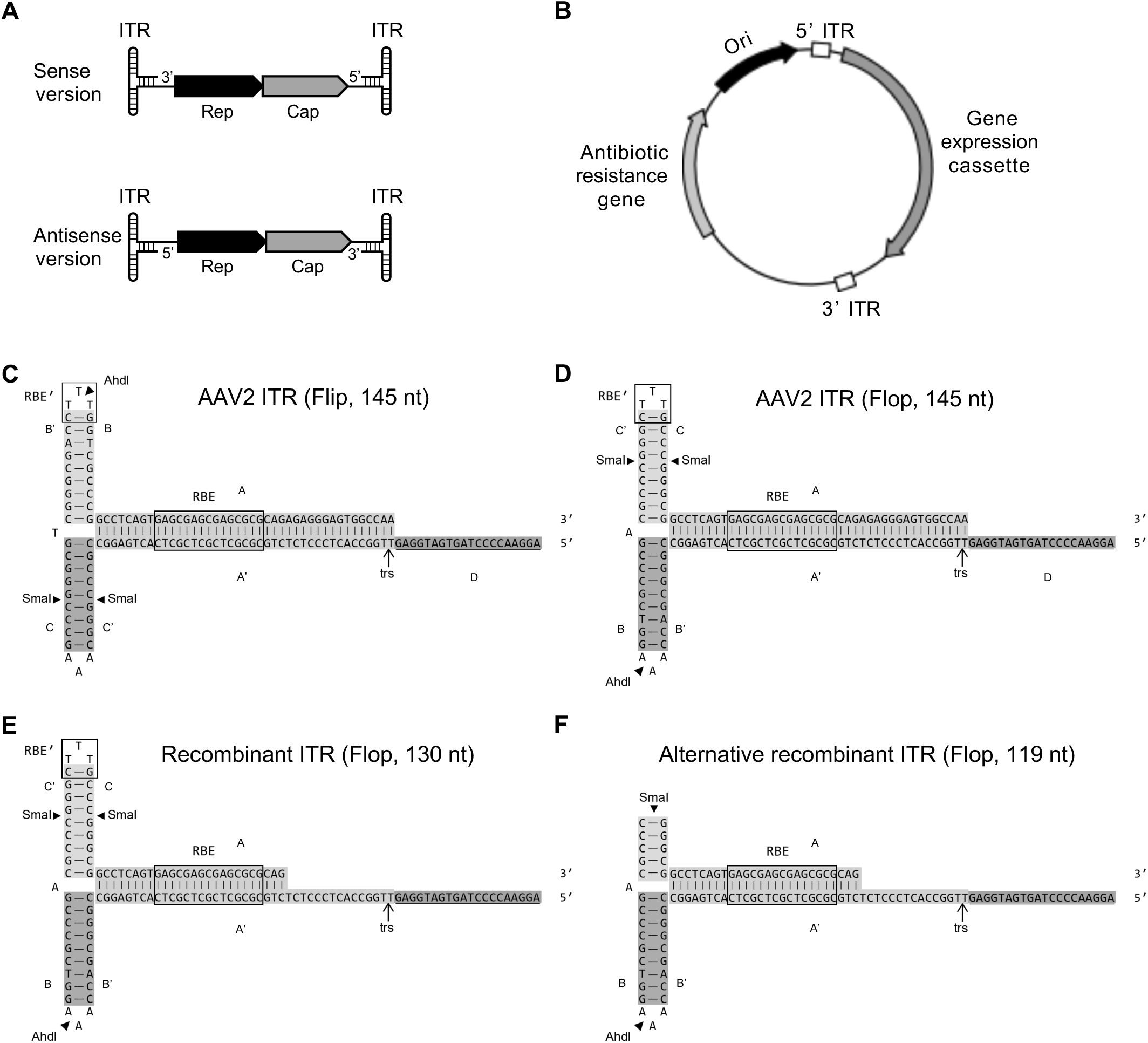
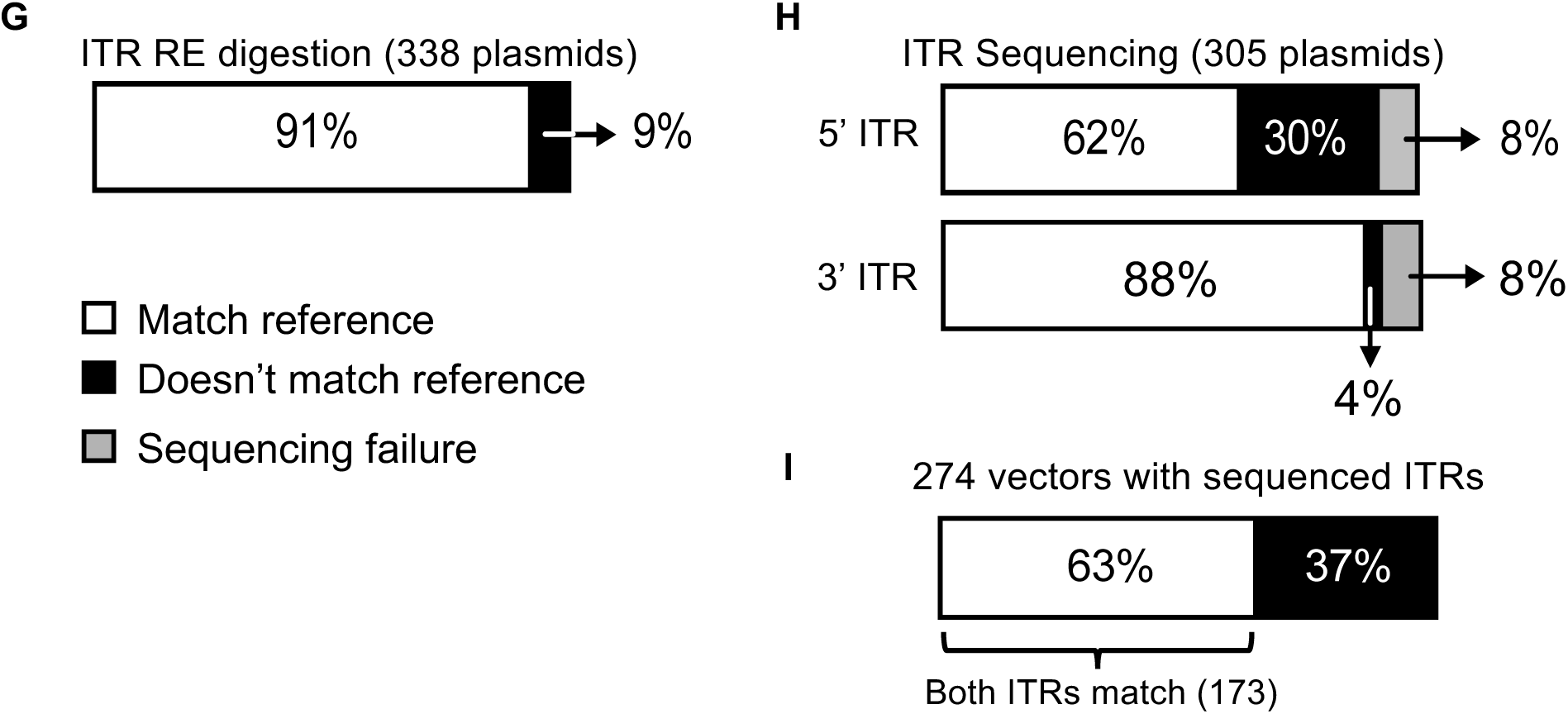
High mutability of ITRs in AAV transfer plasmids. The AAV2 viral genome is a ∼4.7 kb, single-stranded DNA containing Rep and Cap genes flanked by two ITRs on its ends that reverse complement each other (**A**).The typical recombinant AAV transfer plasmid contains two ITR regions flanking the expression cassette (**B**). The secondary structure of the 145-nt AAV2 viral genome ITR is shown in either flip (**C**) or flop (**D**) configuration. It contains three self-annealing regions, A-A’, B-B’, and C-C’, and a single-stranded extension, D, before reaching the internal sequence of the viral genome. The A-A’ stem contains the Rep-binding element (RBE) and the terminal resolution site (trs). The C-C’ arm contains a secondary Rep-binding element (RBE’). The ITRs of AAV transfer plasmids typically have three versions. One is 145 bp long that corresponds to the flop version of ITR in the AAV2 viral genome (**D**). The second is 130 bp long and corresponds to the AAV2 viral genome ITR minus the terminal 15 nt (**E**). The third has an additional 11-bp deletion and is 119 bp long (**F**). The integrity of B-B’ or C-C’ arms on recombinant AAV transfer plasmids can be assayed by restriction enzyme (RE) digestion using *AhdI* or *SmaI*, respectively. (**G**) ITR integrity of 338 AAV transfer plasmids as assayed by RE digestion. (**H**) Integrity of either 5’ or 3’ ITR in the 304 AAV plasmids with correct RE digestion, as further assayed by Sanger sequencing. (**I**) Combined integrity of 5’ and 3’ ITRs in the 274 plasmids for which both ITRs were sequenced.

The wildtype AAV2 viral genome is single-stranded DNA of ∼4.7 kb and can exist as either the sense or antisense strand relative to the direction of the encoded genes Rep and Cap (Figure 2A). The two ITRs that bookend the AAV2 viral genome reverse complement each other (namely, one ITR in an AAV2 genome is identical in sequence to the reverse complement of the other ITR). Each ITR is 145 nucleotides (nt) long that includes a 125-nt self-annealing sequence forming a T-shaped hairpin with two arms denoted B-B’ and C-C’, and a stem A-A’, as well as a 20-nt single-stranded D region that extends from the hairpin (Figure 2C, 2D). The A-A’ stem of ITR contains the Rep-binding element (RBE) and the terminal resolution site (trs). RBE is necessary for recruiting the Rep proteins that replicate the viral genome, while trs is used as the replication initiation site (12). There is also a secondary Rep-binding element (RBE’) on the C-C’ arm that contributes to Rep recruitment (13). The relative positioning of B-B’, C-C’, and A-A’ regions determines whether the configuration of an AAV2 viral genome’s ITRs is “flip” (Figure 2C) or “flop” (Figure 2D), with the former having the B-B’ region, while latter having C-C’, closest to the open end of the AAV2 genome. For either flip or flop configuration, the ITR can exist as the strand with a 5’ open end or the strand with a 3’ open end. For simplicity, only the latter forms are depicted in detail in Figure 2.

The ITRs in AAV transfer plasmids almost all correspond to the flop configuration. We noticed three versions. One is a 145-bp sequence that is the same as the flop version of the full-length ITR sequence in the AAV2 viral genome as depicted in Figure 2D. But this version is very rarely used, with just a few examples out of the hundreds of AAV plasmids that we came across. The second version, which is the most prevalent, is a 130-bp sequence that corresponds to the first version except missing 15 bp at the end of the A region (Figure 2E). When transfer plasmids carrying this ITR are packaged into virus, the missing sequence is added back to form the complete viral genome ITR by copying from the A’ region. Transfer plasmids for which both ITRs are the 145-bp or 130-bp version can generate comparable amounts of AAV viral particles with similar transduction capability (11). These two versions are therefore referred as the wildtype, with one being full-length and the other partial. The third version is a 119-bp sequence that corresponds to the second version but with an additional 11-bp deletion encompassing RBE’ in the C-C’ region (Figure 2F). It is referred to the 119-bp deleted ITR. As discussed later, AAV plasmids carrying one wildtype and one deleted ITR, but not both deleted ITRs, can still be packaged into virus. Note that Figure 2E and 2F depict single-stranded DNA secondary structure based on the sense strand of the ITR sequence on the AAV transfer plasmid, rather than the actual ITR sequence in the recombinant AAV genome being produced upon virus packaging.

To comprehensively assess the fidelity of ITRs on AAV transfer plasmids, we analyzed 338 AAV transfer plasmids (including 2 self-complementary AAV) sourced from academic and industrial labs worldwide. We first subjected them to RE digestion using either *SmaI* or *AdhI*, two enzymes with recognition sites in both wildtype and 119-bp deleted versions of ITRs as depicted in Figure 2, along with one or two enzymes that cut at sites away from the ITRs. This detects mutations in ITRs that abolish *SmaI* and/or *AdhI* cut sites. The assay revealed that about 9% of the plasmids (29/338) had inconsistent patterns compared to that predicted from the senders’ reference ITR sequences (Figure 2G). The 5’ and 3’ ITRs of the 305 RE-validated AAV plasmids were subjected to Sanger sequencing, which revealed that approximately 30% (92/305) of the 5’ ITRs carried mutations relative to their reference sequences (Figure 2H). Interestingly, 3’ ITRs seemed more stable, with only around 4% (13/305) of the plasmids showing mutations relative to reference. Additionally, Sanger sequencing failed for 8% of the 5’ and 3’ ITRs (Figure 2H), presumably due to their high GC content and complex secondary structure that, in the context of some plasmids, are recalcitrant to Sanger sequencing. Since such failed sequences would also be challenging for NGS and they only make up a tiny portion of our plasmids, we did not try to resolve them, and this should not change the conclusion we draw. Of the 274 AAV plasmids with both ITRs successfully sequenced, only 63% (173/274) had both ITR sequences consistent with the senders’ reference (Figure 2I). All counted, about 40% of all the surveyed AAV transfer plasmids had at least one ITR deviating from the wildtype sequence. These results revealed the alarming instability of ITRs in AAV plasmids, especially the 5’ ITR.

We also analyzed the sequences of AAV transfer plasmids deposited at Addgene. We specifically chose the most popular ones that have been requested for more than 100 times and given a blue flame status that stands for the highest tier of popularity. There were 49 blue-flame plasmids with complete depositors’ and Addgene generated whole plasmid sequences (data collected in January 2025). By comparing two versions of the sequences, about 69% (34 out of 49) of the plasmids had mutations in their 5’ ITR and no mutation in 3’ ITR (Figure 4A, Table S4). These results further supported the stunningly high instability of ITRs and the interesting phenomenon that 5’ ITR displays much greater instability than the 3’ ITR. ITR sequences of these blue-flame plasmids, except for one self-complementary AAV (scAAV) plasmid, are included in further analysis below.

#### Stability of ITRs in AAV transfer plasmids is affected by flanking sequences

We examined whether sequences immediately flanking the ITRs on the AAV transfer plasmids would impact their stability. Based on reference 5’ ITR sequences from the senders, 274 plasmids were classified into four distinct groups distinguished by the 5’ ITR sequence itself and the nature of its upstream flanking sequence (Figure 3). Group A consists of 147 plasmids whose sender-provided 5’ ITR reference sequences matched the 130-bp wildtype ITR version shown in Figure 2E, and additionally, the flanking sequence immediately upstream of the 5’ ITR contained an 11-bp high-GC (73%) sequence (Figure 3A). Upon Sanger sequencing, we found that around 61% (90/147) of the Group A plasmids contained mutations in their 5’ ITRs relative to the reference (Figure 3A). Among the mutations, the most prevalent, which occurred in 87 out of 90 cases, was the deletion of 11 bp in the C-C’ region (Figure 3A, Table S5), which effectively converted the 5’ ITR from the 130-bp wildtype version shown in Figure 2E into the 119-bp deleted version shown in Figure 2F. Additionally, 5’ ITRs on two plasmids exhibited a 22-bp deletion, and on one plasmid, a 4-bp deletion (Figure S1). Moreover, 29 of the 48 blue-flame ssAAV transfer vectors from Addgene share the same high-GC flanking sequences of Group A plasmids. Compared to their depositors’ reference maps, Addgene sequencing revealed that 28 of these contained the same 11-bp deletion in the C-C’ region while no mutation in the 3’ ITR was identified (Figure 4B, Table S4). In a single study (14), the same 11-bp deletion happened to the 5’ ITR of 4 plasmids (#60226, 60227, 60229, and 60231) out of the 7 analyzed (#60225-60231). Collectively, these results showed that the 5’ ITR with high GC content in the flanking region is very instable.

**Figure 3.**
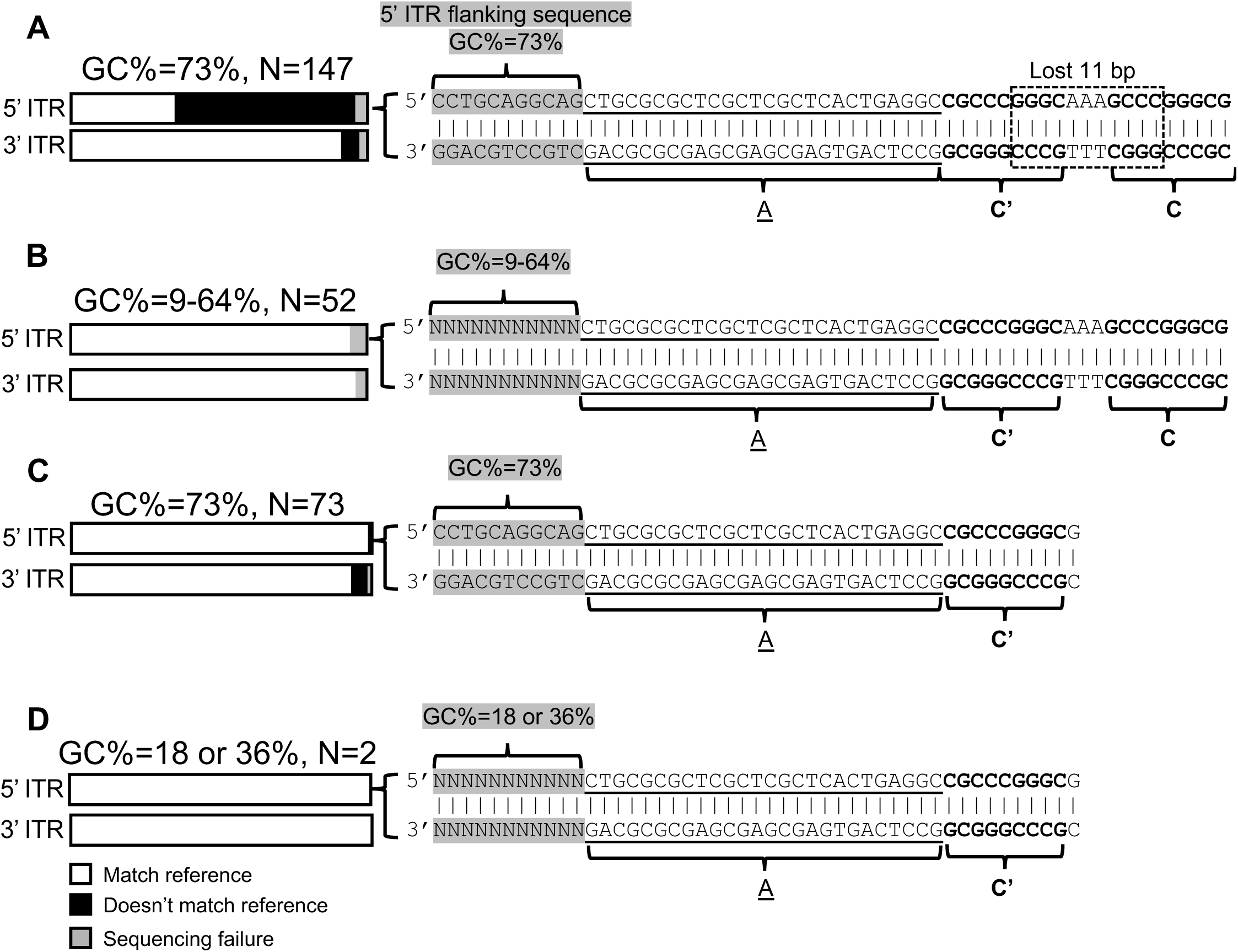
Correlation between ITR stability and GC content in the 11-bp flanking sequence of three groups of 5’ ITR. (**A**) 5’ ITR in 147 plasmids have a high GC content (73%) 11-bp flanking sequence, and the sequence of 90 5’ ITR (∼61%) were unmatched with the user-provided reference. 87 of the unmatched 5’ ITR lost the 11 bp in the C arm (boxed with dash line) compared to their reference. (**B**) 52 plasmids have the exact same 5’ ITR as (**A**) but 9-64% GC content in the 11-bp flanking sequence. No mutation was detected in the 5’ ITR of these plasmids. (**C**) 73 plasmids had the exact same high-GC 11-bp flanking sequence but the alternative 5’ ITR sequence missing the 11 bp in the C arm. The 5’ ITR sequence of one plasmid unmatched its reference. (**D**) 2 plasmids had the alternative 119-bp 5’ ITR but different flanking 11 bp sequence of lower GC content.

**Figure 4.**
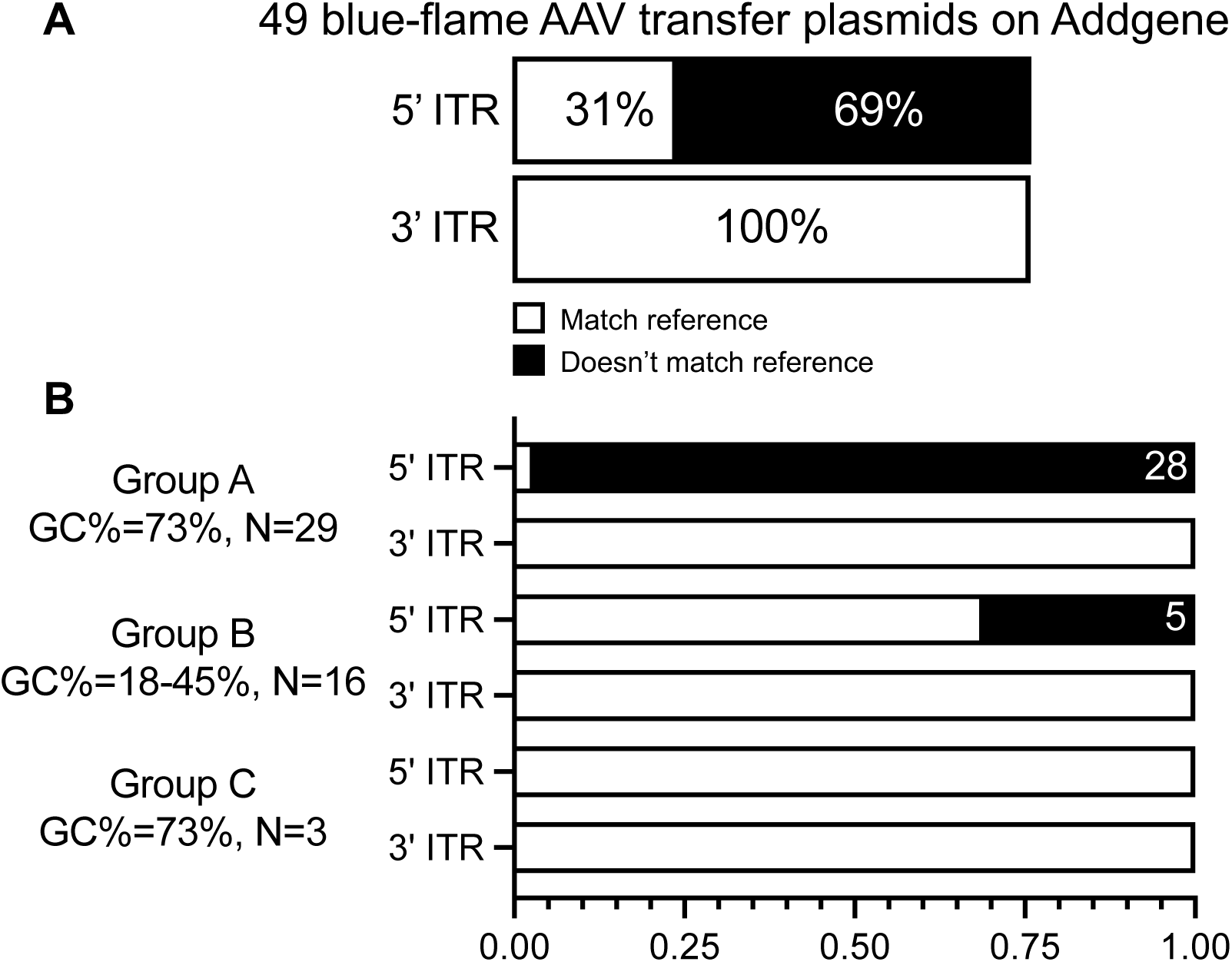
Mutability of ITRs in AAV transfer plasmids with blue flame on Addgene. (**A**) The depositors’ and Addgene generated whole plasmid maps of 49 blue-flame AAV transfer plasmids were aligned to check the mutations in ITRs. (**B**) 48 plasmids were categorized into three groups based on the same features shown in Figure 3A-C. Addgene’s sequencing showed that 5’ ITR in 28 out of 29 plasmids with a high GC content (73%) 11-bp flanking sequence (Group A) do not match their depositor’s maps. All mutations were the lost of 11 bp highlighted in Figure 3A. For 16 plasmids with lower GC content (18-45%) in the 5’ ITR flanking sequence (Group B), 5 of their 5’ ITR sequences do not match with the depositor’s. No mismatches in Group C plasmids and all 3’ ITRs were observed.

Group B consists of 52 plasmids whose sender-provided 5’ ITR reference sequence also matched the 130-bp wildtype ITR version shown in Figure 2E, but the flanking sequence immediately upstream of the 5’ ITR had GC content ranging from 9% to 64% (Figure 3B, Table S5). Strikingly, none of the plasmid in this group had a different 5’ ITR from its reference except for 3 plasmids whose 5’ ITR was not fully sequenced. There were 16 blue-flame AAV plasmids belonging to Group B, and their GC content in the 5’ ITR flanking region ranges from 18-45%. Of these, 5 had mutations in 5’ ITR based on Addgene’s sequencing (Figure 4B, Table S4).

Group C consists of 73 plasmids whose sender-provided 5’ ITR reference sequence matches the 119-bp deleted version as depicted in Figure 2F, and which also have the same 11-bp high-GC (73%) flanking sequence immediately upstream of the 5’ ITR as found in Group A (Figure 3C). Among them, only one contained a mutation that is a 15-bp deletion (Figure S2, Table S5). Lastly, Group D consists of 2 plasmids with the identical 119-bp 5’ ITR as the Group C plasmids, yet the flanking 11 bp contain 18 or 36% GC (Figure 3D), and none are mutated. Only 3 out of 48 blue-flame AAV plasmids from Addgene belong to Group C, and no mutation was found in 5’ ITRs (Figure 4B, Table S4).

Thus, the coupling of the wildtype version of 5’ ITR with high-GC flanking sequence appears to lead to greatly increased mutability, with the 11-bp deletion in the C-C’ region being the most prevalent mutation. This hypothesis aligns with previous finding of ITR instability on AAV plasmids when flanked by a 15-bp sequence of 100% GC, which was markedly improved when this flanking sequence was eliminated (9). It also aligns with our own experience that during cloning, using AAV plasmid backbones in which ITRs are flanked by high-GC sequences tends to produce more clones bearing mutations in the ITRs. Indeed, when we cultured *E. coli* carrying a plasmid with two validated 130-bp ITRs flanked by the 11-bp high-GC sequence as shown in Figure 3A for ten passages, we found that about half of the plasmid DNA now carried mutated ITR, indicating a remarkably high mutation rate (unpublished data). Further supporting our hypothesis, our recent data showed that reducing the GC content in the sequence flanking the ITR and restructuring the functional elements on the plasmid backbone can significantly increase ITR stability (manuscript in preparation).

The above observations also argue that the 119-bp deleted version of 5’ ITR in Figure 2F, even when annotated in the reference sequence as such, was not created intentionally for a purpose by someone, but actually resulted from frequent deletions occurring to the 130-bp wildtype ITR in Figure 2E when flanked by the 11-bp high-GC sequence that caused instability to the ITR. This prevalent mutation probably occurred independently in multiple labs, which then likely passed onto many other labs. It is possible that some researchers were unaware of this mutation having occurred in their AAV plasmids and still assumed the wildtype 5’ ITR sequence as the reference, while some other researchers saw the mutated sequence at some point and just considered it to be the correct reference. The same 119-bp deleted 5’ ITR were also observed in two plasmids for which the 11-bp upstream flanking sequence contained low GC (18% or 36%). For these, it is possible that the 11-bp deletion occurred spontaneously in their 5’ ITRs even in the context of low-GC upstream flanking sequence, or the deletion first occurred on a different backbone containing the high-GC flanking sequence, and the ITR-to-ITR region was later subcloned into the current vectors.

We next examined the integrity of 3’ ITRs in AAV transfer plasmids. In Group A, which contained 147 plasmids, 143 sender-provided 3’ ITR reference all bore the same 130-bp sequence and the 11-bp high-GC flanking sequence, just like their 5’ ITR reference sequence (Figure 3A). We found that 8 plasmids in this group had point mutations in their 3’ ITR, while none had mutations in their 5’ ITRs. Based on the senders’ reference, the 3’ ITR on two of the remaining four vectors was of the 119-bp deleted version, flanked by the high-GC 11-bp sequence. The 3’ ITR of the last two plasmids was of the 130-bp version, whose 11-bp flanking sequence had 64% GC. These four vectors had no mutations in their 3’ ITR relative to the senders’ reference. In Group C with 73 plasmids, the sender-provided 3’ ITR reference sequence all had the 130-bp wildtype version, with 72 having the high-GC 11-bp flanking sequence and only one having a low-GC (18%) 11-bp flanking sequence. In four of these plasmids, the 3’ ITRs also mutated to the 119-bp deleted version, such that both their ITRs were of the deleted version. No mutations were found in the 3’ ITR of Groups B and D plasmids.

There are two important take-home messages from the above data. First, the same 130-bp ITR sequence with the 11-bp high-GC flanking sequence can be exceedingly mutable when it is the 5’ ITR, and moderately mutable when it is the 3’ ITR (Figure 3). This suggests that there are other factors affecting ITR stability, which we hypothesize to be the distance from the ITR to the plasmid replication origin (Ori). The 5’ ITR is usually 200-500 bp from Ori, whereas the 3’ ITR is typically over 2 kb away from Ori. Second, both ITRs of a transfer plasmid can be mutated, and when this happens, virus packaging is severely impaired as discussed below.

It has been shown that when packaged into virus, AAV transfer plasmids carrying a mutant ITR on one end and a wildtype ITR on the other end can produce viral genome for which the mutant ITR is repaired, presumably by templating off of the wildtype ITR (15). This notwithstanding, how different types of mutant ITRs influence packaging efficiency, viral genome integrity, and intended biological functions of the virus is not well understood. Furthermore, once mutations have occurred to one ITR, such as the 11-bp deletion that converts the 130-bp wildtype version to the 119-bp deleted version, additional mutations can still happen to the other ITR at a reasonable frequency. When both ITRs are mutated, the repair mechanism is no longer effective, and AAV packaging will be seriously compromised (11). Caution is thus advised when using AAV vectors in gene therapy applications where ITR fidelity could impact drug efficacy and safety. We suggest that AAV transfer plasmids whose 5’ and 3’ ITRs are both the wildtype version, and which do not show a strong tendency to mutate (such as the high mutability observed for the ITR with the high-GC flanking sequence), are preferrable over other designs in gene therapy applications.

#### Good vector practice

Recognizing the critical importance of vectors in research and medicine, and the high prevalence of errors in lab-made plasmids, we recommend the following “good vector practice” (GVP) based on our extensive experience with constructing and validating hundreds of thousands of plasmids. We believe that the adoption of GVP by the research community can improve the quality of vectors in labs worldwide.

GVP is a system that covers the whole life cycle of a vector, namely vector design, construction, in-process QC, final QC, banking and management. To design a potent and stable vector, researchers should carefully consider key factors including but not limited to the choice of vector system (e.g., viral, transposon, or transient transfection vector), backbone type (e.g., high-copy, low-copy, or miniaturized), antibiotic selection marker (e.g., ampicillin or kanamycin), essential biological components (e.g., promoters, linkers, spacers, polyadenylation signals, and enhancers), and codon optimization of coding sequences for enhanced expression, stability, and cloning efficiency. Since even experienced molecular biologists may overlook certain nuances, it is important to avoid the commonly observed design errors listed in Table 1, and if needed, consult experts from leading resources like VectorBuilder, iGEM, and Addgene. Another common challenge is choosing well-documented and validated versions of components. For instance, multiple versions of CMV promoter with various length and sequences have been reported. We refer researchers to our Vector Design Studio, a computer-aided vector design tool available online (https://www.vectorbuilder.com/design.html), which has incorporated a comprehensive set of validated design algorithms capable of creating optimized vector designs from standardized and validated vector systems and components. Additionally, our upcoming Gene Delivery Forum described above will be dedicated to supporting the community by promoting knowledge-sharing regarding vector design and the standardization and cross-referencing of vector systems and components. Finally, the optimization of vector design should be seen as an iterative process guided by experimental results.

When initiating a plasmid cloning project, it is essential to verify the fidelity of DNAs used as starting materials, especially if they are from sources that don’t routinely perform stringent QC such as lab peers and collaborators. Assays such as diagnostic restriction digestion, sequencing and other adequate QC tests should be performed as needed to ensure that the starting DNA materials are consistent with their theoretical sequence and structure.

In-process QC should be considered for projects involving multistep cloning. In particular, all steps involving DNA synthesis should be regarded as a potential source of mutations, such as reverse transcription, PCR, and Gibson assembly, and these steps should employ proper methods to mitigate the risk of mutations (e.g., using high-fidelity polymerase) and should pass QC (i.e. sequence validation) before proceeding.

Final QC should be done for more than just the inserted sequences, but also preexisting components on vectors that are vital for downstream applications. Sequences on viral vectors that function in packaging but not downstream biological applications, which tend to get overlooked, should be confirmed to ensure proper viral packaging. This is especially the case for AAV ITRs, which tend to mutate.

A free-standing vector banking and management system apart from regular lab reagent storage and note-keeping is essential for maintaining high fidelity and traceability of lab vectors over the long run. Plasmids in a given lab, whether sourced or self-built, should have unique IDs, descriptive names that reflect their purpose and key components, and complete records including creators, maps, sequences, cloning methods and steps employed, and QC results.

Proper archiving of this information is critical for troubleshooting any issues that may arise in downstream applications. For plasmids retrieved from frozen stocks that have not been used for a longer time, it is important to re-validate them to ensure consistency with past records.

As a key aspect of the academic ecosystem to maintain vector quality, we suggest that journals encourage or even require authors to utilize GVP principles when publishing. This would significantly raise vector integrity in research and medicine and will improve the reproducibility of molecular data.

The GVP concept is especially important to vectors intended for clinical use where additional considerations in vector safety, efficacy and manufacturability must be addressed. When designing a clinical-grade vector, safety is paramount, and certain components commonly used in research vectors may need to be substituted to meet regulatory requirements on safety in human use. For instance, due to the allergenic potential of residual ampicillin, the frequently used ampicillin resistance marker on plasmid backbones should be replaced with a safer alternative such as kanamycin resistance or antibiotic-free backbones. Similarly, the WPRE (woodchuck hepatitis virus post-transcriptional regulatory element), often used to enhance expression, may need to be substituted with safer varieties (e.g., WPREmut6 or WPRE3) due to its potential oncogenic risk. To achieve optimal efficacy, clinical-grade vectors should be more comprehensively optimized at the design stage than research-grade vectors, with careful selection of backbones, coding sequences, promoters, and regulatory elements. Furthermore, the manufacturability of clinical-grade vectors in terms of yield, long-term sequence stability, and the percentage of supercoiled plasmid, can significantly impact the financial viability of large-scale commercial production. We have encountered many cases where safety and manufacturability issues of vectors were not addressed during the preclinical stage of gene therapy development pipelines because attention was solely given to efficacy, only for these issues to surface during GMP production when the vectors enter clinical trials. These projects had to be paused to redesign the vectors, such as changing plasmid backbones, adjusting GC content, reducing repetitive sequences, etc., which can cause major setbacks in both time and money, and in some cases, even lead to complete redoes of preclinical testing of the vectors. In the world of drug development, this can mean the life and death of a multimillion-dollar program. To prevent such costly mistakes, we regard vector designs that incorporate safety requirements and the subsequent testing of manufacturability, preferably on several candidate designs, as a crucial and routine aspect of QC in clinical vector development.

## Discussion

For many decades, researchers have made their own customized plasmids in the lab to meet their specific research needs. Yet, not only can lab-made plasmids be constructed with mistakes, they are also subject to ongoing mutations while maintained in *E. coli* due to DNA replication errors (16), homologous recombination (17), and insertions of jumping sequence elements (18). It is therefore surprising that despite the ubiquity and critical importance of lab-made plasmids in research and medicine, there is as yet no systematic assessment of their quality.

Being a cloning service provider, we had the opportunity to handle a large number of plasmids sourced from academic and industrial labs around the world. We report, for the first time, a large-scale quality assessment of lab-made plasmids, which showed, much to our surprise, a very high rate of errors. We found that approximately 15% of plasmids had design errors of varying significance, and about 35% contained sequence errors in functional regions (excluding AAV ITRs) (Figure 1). For AAV transfer plasmids, about 40% had mutations in their ITRs relative to the wildtype form. In total, we estimate that 45-50% of lab-made plasmids have undetected design and/or sequence errors that could potentially compromise the intended applications. We did not observe any significant bias in the prevalence of errors based on researcher demographics, the nature of their affiliation (e.g., academic vs. industry), or the intended purpose of the plasmids. Indeed, we suspect that this number may underestimate the true scale of quality issues in lab-made plasmids because we had asked our clients to check the designs and sequences of their plasmids before submission to us, and also because they were paying for our services utilizing their plasmids. We also encourage other cloning service providers, including both commercial companies and academic core facilities, to report the errors in lab-made plasmids that they find. Such collective efforts can raise community awareness about this issue, thereby improving the trustworthiness of scientific discoveries and therapeutic products made with this essential molecular biology tool.

The high error rate of lab-made plasmids suggests that many researchers building their own vectors may lack the sophisticated and nuanced expertise needed to properly design vectors, or are under time and cost constraint to conduct adequate quality control of the vectors constructed or propagated in their labs. Even when the principal investigator and experienced members of the lab have adequate expertise in vector construction and quality control, less experienced members of the lab may not benefit from such expertise due to insufficient training. Our study thus shines a light on the need to include GVP as an important part of lab training.

The issues identified here are even more critical in the development of therapeutics. Based on our clinical-grade plasmid manufacturing experience, we have seen that many drug developers overlook the importance of verifying the design and sequence of their vectors during the R&D phase. As a result, substantial time and resources are invested in suboptimal or even incorrect plasmids, often until just before applying for a clinical trial. At this stage, heightened scrutiny reveals these errors, requiring costly and time-consuming corrections that may necessitate redesigning the plasmid, repeating efficacy and toxicology studies, and redeveloping the clinical-grade manufacturing process, which inevitably leads to significant delays and financial burdens.

Public plasmid repositories, which have greatly facilitated the sharing of published plasmids, often perform independent sequencing of their plasmids. However, these repositories typically do not make statements about whether their plasmids are properly designed. They also typically do not address the sequence stability of plasmids over the course of their possession or the source of any discrepancies between their sequencing data and the depositor-supplied reference sequences. Analysis for the most popular AAV transfer plasmids from Addgene revealed the same worrying trend that we found. Namely, the 5’ ITR is rather unstable. For instance, we observed that Addgene’s sequencing data for their plasmids #112159, #112172, #112173 and #112174, which are AAV vectors from the same researchers, is grossly different in the ITR relative to depositor-supplied references, and the backbone of #112159 and #112173 also appear flipped. Yet, another vector from the same depositor and same study, #112168, has identical ITR sequences between Addgene-generated sequencing data and depositor-supplied reference. The fidelity of plasmids from another public repository, the Registry of Standard Biological Parts, which is a platform for researchers to share biological parts for building synthetic biology systems, also showed concerning issues. A study in 2008 surveyed 1,536 plasmids from the Registry by PCR-amplifying the biological parts and found that about 26% of the reactions generated no or nonspecific amplicons, for which the study did not rule out the possibility of suboptimal PCR conditions (19). Furthermore, the study sequenced and assembled 354 parts from the plasmids, and 20 could not be aligned with published sequences.

Our finding mirrors other reports of major problems with widely used lab reagents that have gone “under-the-radar” for many years simply because researchers did not think to question their quality, such as mycoplasma contamination and misidentified mammalian cell lines (3-5).

We argue that there is a compelling need for community-wide standards and resources to uphold the quality of gene delivery vectors in research and medicine. These may include educational materials on how to design appropriate vectors for various applications, best practices in the construction, propagation, storage, transfer and QC of plasmids and related reagents such as libraries and packaged viruses, and mechanisms that encourage researchers to share their expertise especially tips for improving vector performance and avoiding pitfalls.

## Data availability

All survey results are presented.

## Acknowledgements

This study was supported by VectorBuilder.

---

**Table S1.** Genes with moderate toxicity to E. coli host.

| Gene Name | Species | NCBI Gene ID | Function |
| --- | --- | --- | --- |
| ABCB4 | Human | 5244 | Maintains the cell membrane and recruits phospholipids |
| Actn1 | Mouse | 109711 | Binds actin filaments to the cell membrane |
| Adam10 | Mouse | 11487 | Regulates of shedding of ectodomain of proteins including cell adhesion proteins |
| ADAM17 | Human | 6868 | Regulates of shedding of ectodomain of proteins including cell adhesion proteins |
| Ano5 | Mouse | 233246 | Chloride channel involved in membrane maintenance and repair |
| Cdh2 | Mouse | 12558 | Regulates neural stem cell quiescence by mediating cell adhesion |
| CDK12 | Human | 51755 | Maintains genomic stability and regulates DNA repair |
| CFTR | Human | 1080 | Chloride channel involved in fluid homeostasis |
| Chek1 | Mouse | 12649 | Enhances chromatin assembly and aids in DNA damage repair |
| COL11A1 | Human | 1301 | Collagen responsible for bone development and extracellular matrix contribution |
| DYRK1A | Human | 1859 | Kinase involved in DNA damage repair |
| EP300 | Human | 2033 | Regulates cell proliferation and differentiation |
| GPAT3 | Human | 84803 | Protects against lipotoxicity |
| Gpr5 | Mouse | 620246 | G protein-coupled receptor involved in calcium regulation |
| Gtf3c1 | Mouse | 233863 | Involved in RNA polymerase III-mediated transcription of 5S rRNA and other RNAs |
| HDAC2 | Human | 3066 | Regulates cell cycle progression |
| Htra3 | Mouse | 78558 | Inhibits TGF-β signaling and may act as tumor suppressor |
| ITGA6 | Human | 3655 | An integrin functioning in cell surface adhesion and inhibition of HER2 signaling |
| ITGB8 | Human | 3696 | An integrin functioning in cell surface adhesion, signaling, and vasculogenesis |
| JAK2 | Human | 3717 | Tyrosine kinase involved in cell growth, development, and histone modifications |
| Jmy | Mouse | 57748 | Regulates apoptosis |
| KCNH8 | Human | 131096 | Voltage-gated calcium channel promoting rectifying current |
| KIT | Human | 3815 | Regulates cell survival and proliferation |
| KL | Human | 9365 | Regulates calcium and phosphorous homeostasis via inhibition of vitamin D synthesis |
| KLHL31 | Human | 401265 | Inhibits stress-activated JNK pathway and promotes apoptosis |
| LMNB1 | Human | 4001 | Maintains nuclear envelope and may interact with chromatin |
| LRRK2 | Human | 120892 | Involved in retrograde trafficking pathway of recycled proteins |
| Nfkb2 | Mouse | 18034 | Activates genes involved in inflammation and immune response |
| nompC | Drosophila | 33768 | Calcium channel involved in sensing mechanical stimuli |
| NPC1 | Human | 4864 | Involved in cholesterol processing |
| PAK1 | Human | 5058 | Kinase involved in cytoskeletal dynamics, proliferation, and apoptosis |
| PARP2 | Human | 10038 | Recruits DNA repair factors following DNA damage |
| PIEZO2 | Human | 63895 | Calcium channel involved in sensing mechanical stimuli |
| Piezo2 | Mouse | 667742 | Calcium channel involved in sensing mechanical stimuli |
| PIK3CA | Human | 5290 | Regulates pathways involved in cell growth, survival, and proliferation |
| PRKN | Human | 5071 | Regulates processing of damaged mitochondria and inhibits apoptosis |
| SCN10A | Human | 6336 | Sodium-selective channel that mediates ion permeability across excitatory membranes |
| Scn2a | Rat | 24766 | Sodium-selective channel that mediates ion permeability across excitatory membranes |
| SCN4A | Human | 6329 | Voltage-gated sodium channel subunit necessary for initiation of action potentials |
| Set | Mouse | 56086 | Inhibits nucleosome acetylation and apoptosis |
| SLC38A9 | Human | 153129 | Activates TOR pathway members to promote cell growth |
| SMARCA1 | Human | 6594 | Involved in chromatin remodeling and regulation of apoptosis |
| Tjp2 | Mouse | 21873 | Involved in tight junction and adherens junction formation |
| TRPA1 | Human | 8989 | Calcium channel that serves as chemical and mechanical stress sensor |
| TRPS1 | Human | 7227 | Transcriptional repressor that regulates proliferation |
| WRN | Human | 7486 | Maintains genomic stability and regulates DNA repair |

**Table S2.** Genes with severe toxicity to E. coli host.

| Gene Name | Species | NCBI Gene ID | Function |
| --- | --- | --- | --- |
| Abcb1a | Mouse | 18671 | Transporter involved in multi-drug resistance |
| Abcb1a | Rat | 170913 | Transporter involved in multi-drug resistance |
| APOBEC3B | Human | 9582 | Deaminates single stranded viral DNA |
| AT5G17850 | Thale Cress | 831653 | Sodium/calcium exchange transporter involved in regulating intracellular calcium levels |
| cac | Fruit fly | 32158 | Voltage-gated calcium channel involved in neurotransmitter release |
| CAX7 | Thale Cress | 831654 | Cation/calcium exchange transporter involved in regulating intracellular calcium levels |
| DSG3 | Human | 1830 | Mediates intermediate filament function in cell-cell adhesions |
| GOLIM4 | Human | 27333 | Assists in protein transport through the Golgi apparatus |
| MRC1 | Human | 4360 | Functions in recognition of pathogens |
| Mrc1 | Mouse | 17533 | Functions in recognition of pathogens |
| Nalcn | Mouse | 338370 | Leaky sodium channel that regulates membrane electrical potential |
| NCAPD3 | Human | 23310 | Regulates chromosomal architecture and segregation |
| NF1 | Human | 4763 | Regulates cell growth and differentiation |
| pkd2 | Zebrafish | 432387 | Calcium-activated nonspecific cation channel involved in cilium development |
| PRG4 | Human | 10216 | Growth factor that regulates cell adhesion |
| SCN1A | Human | 6323 | Sodium-selective channel that mediates ion permeability across excitatory membranes |
| SCN2A | Human | 6326 | Voltage-gated sodium channel subunit necessary for initiation of action potentials |
| SCN3A | Human | 6328 | Voltage-gated sodium channel subunit necessary for initiation of action potentials |
| Scn7a | Mouse | 20272 | Sodium-selective channel that mediates ion permeability across excitatory membranes |
| Scn8a | Human | 20273 | Voltage-gated sodium channel subunit necessary for initiation of action potentials |
| SCN9A | Human | 6335 | Voltage-gated sodium channel that plays a role in inflammatory pain |
| SF3B1 | Human | 23451 | Splicing factor that removes introns from pre-mRNAs |
| Slco1b2 | Mouse | 28253 | Solute carrier organic anion transporter involved in bile transport |
| TLR4 | Yangtze finless porpoise | 112411200 | Initiates immune activation in response to pathways associated with damage and pathogens |
| Top1 | Rat | 64550 | Topoisomerase that removes supercoiling by making single strand cut and regulating rejoining |

**Table S3.**
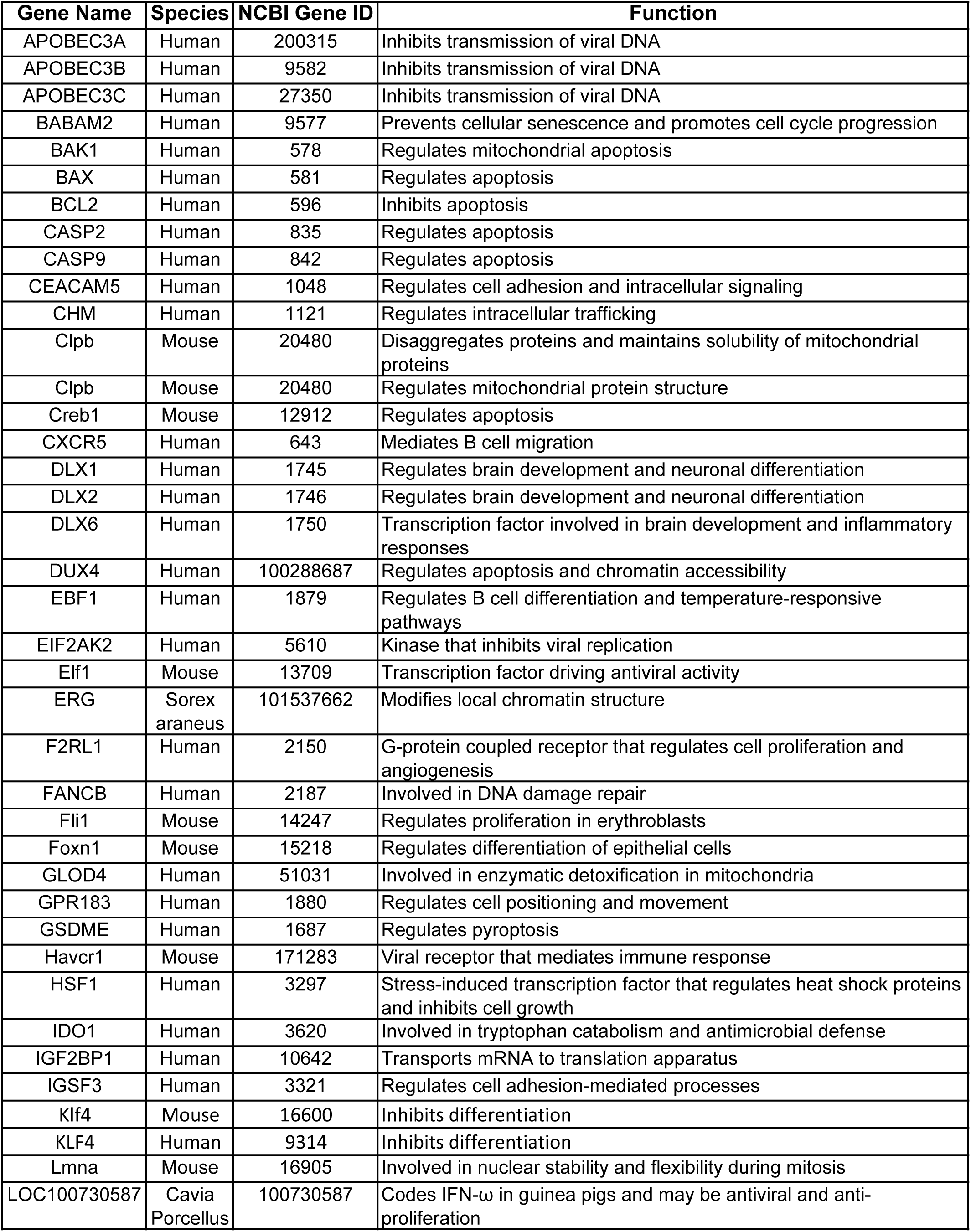
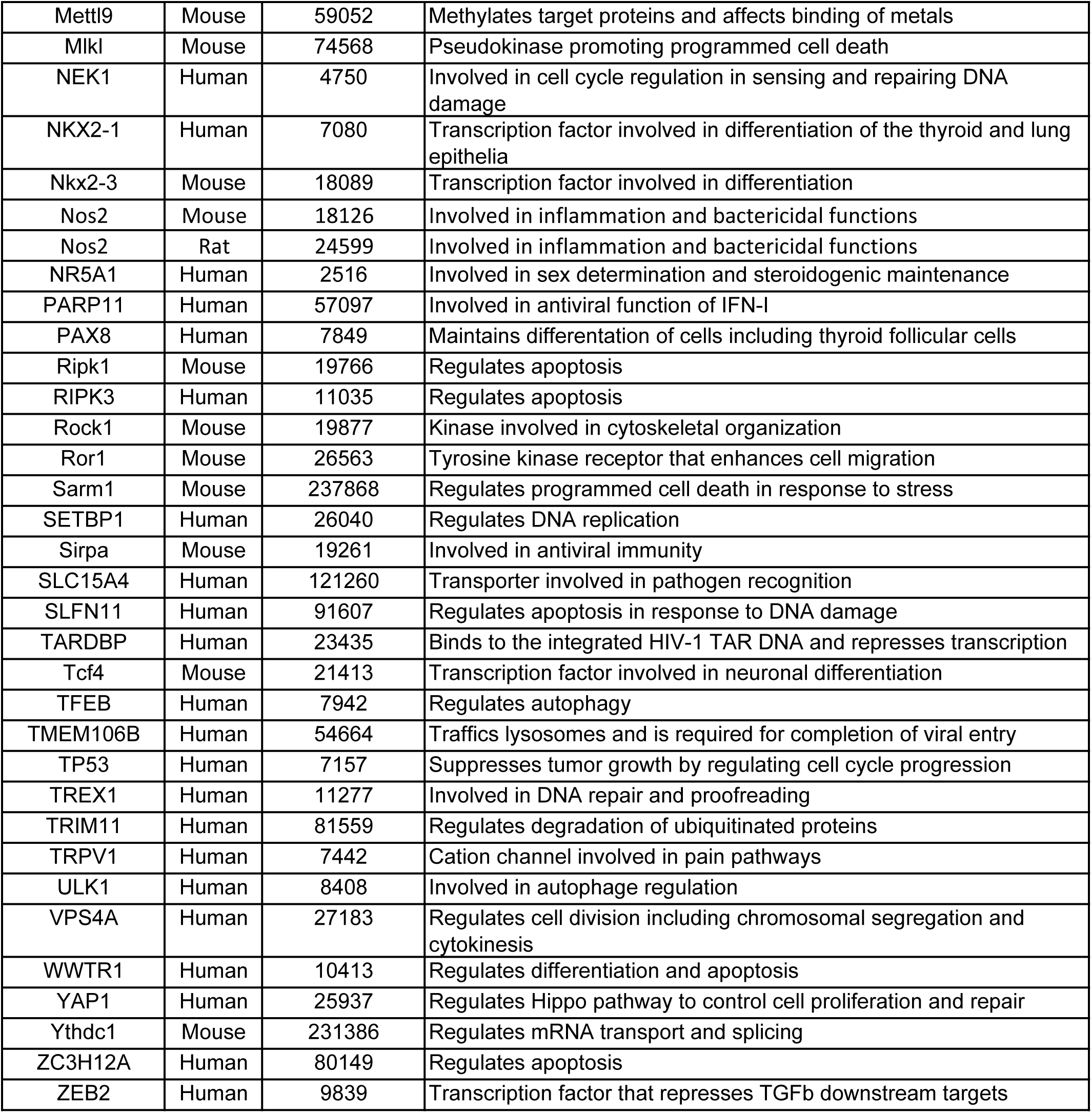
Genes with toxicity to packaging cells.

**Table S4.**
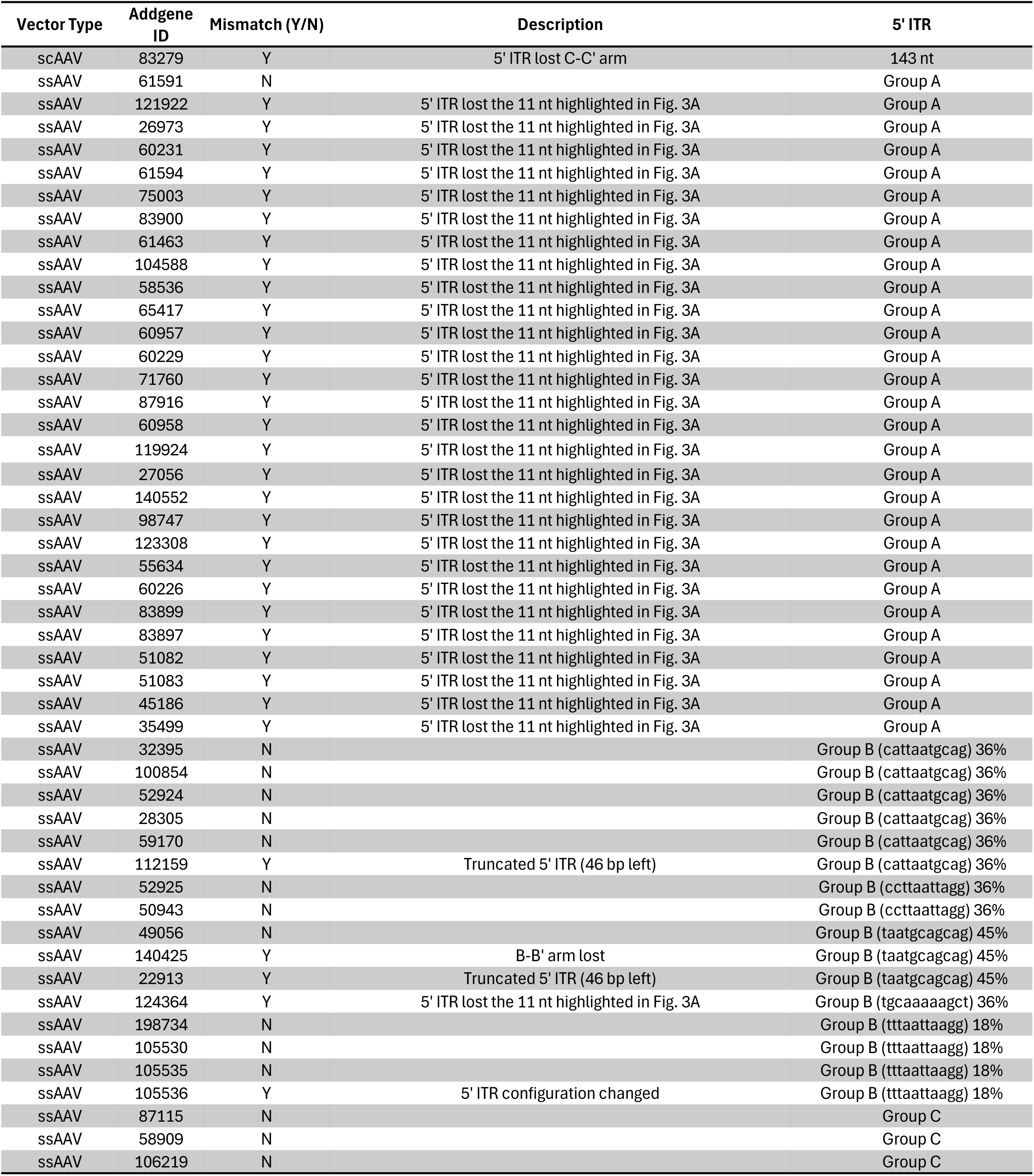
49 blue-flame AAV transfer plasmids with complete Addgene sequenced and depositor’s maps.

**Table S5.**
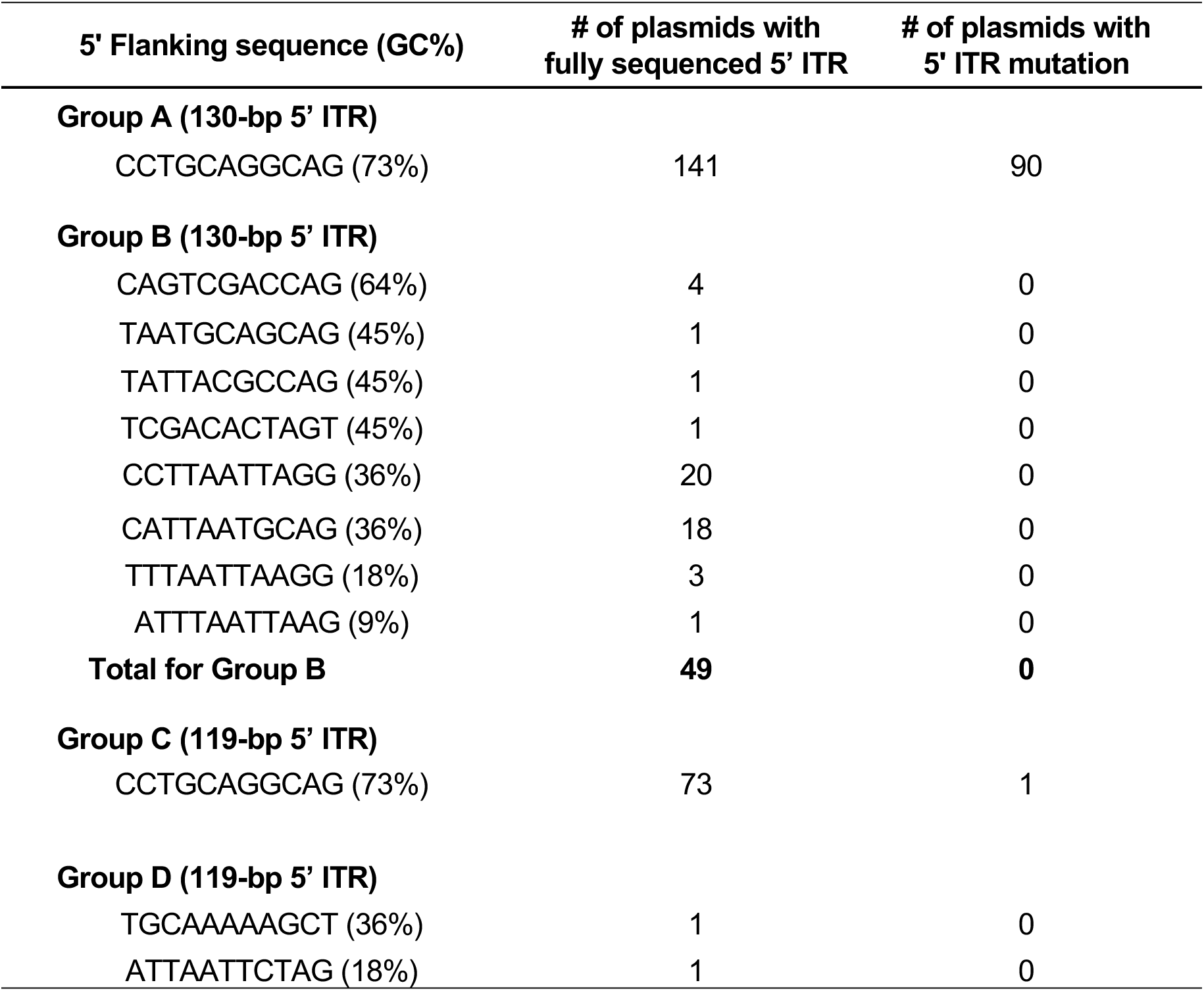
Types of fully sequenced 5’ ITR flanking sequences and their associated 5’ ITR mutations.

**Figure S1.**
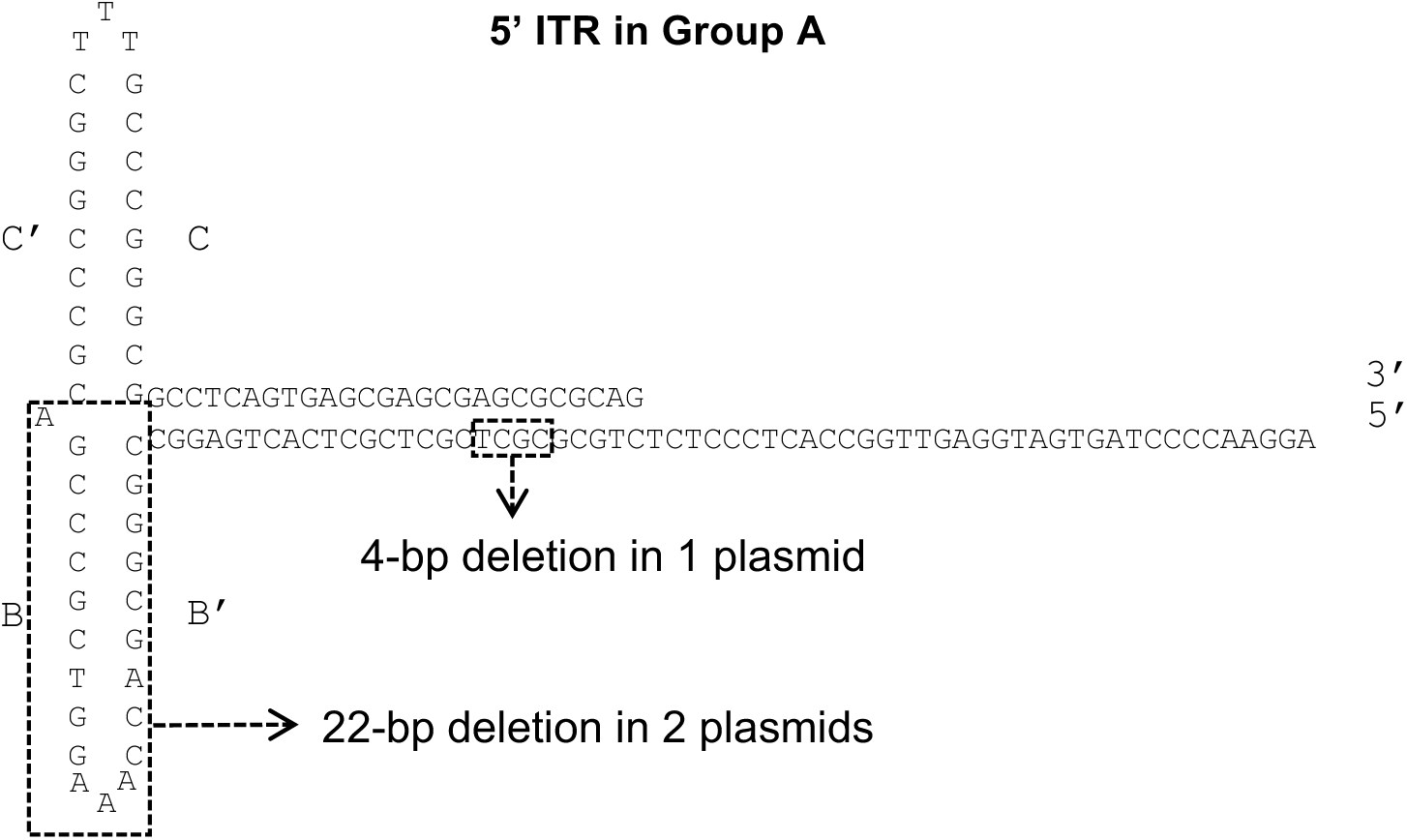
Deletions identified in 5’ ITR of Group A transfer plasmids (Figure 3A) that are not the 11-bp deletion associated with the 119-bp deleted version of ITR.

**Figure S2.**
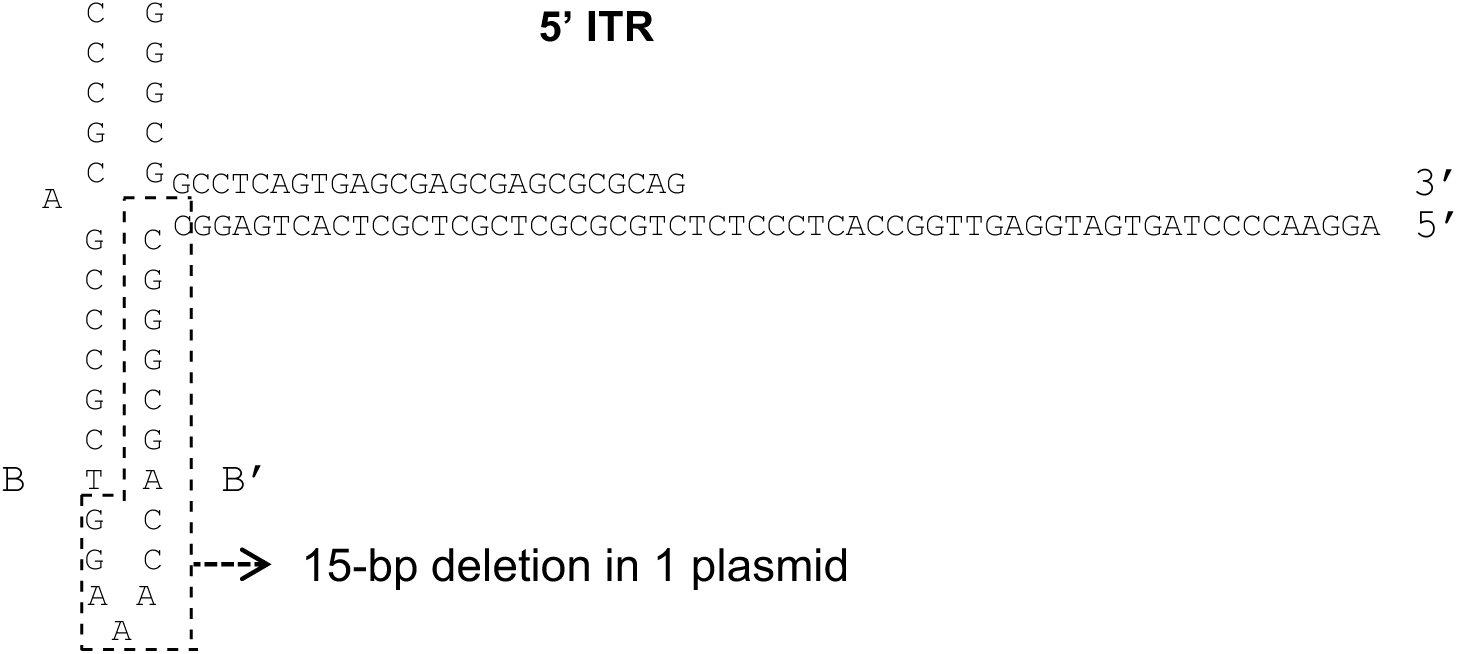
Deletion identified in 5’ ITR of Group C transfer plasmid (Figure 3C).

